# Environmental factors drive bacterial degradation of gastrointestinal mucus

**DOI:** 10.1101/2025.01.16.633397

**Authors:** Sandra L. Arias, Ellen W. van Wijngaarden, Diana Balint, Joshua Jones, Carl C. Crawford, Parul J. Shukla, Meredith Silberstein, Ilana L. Brito

## Abstract

The mucus layer lining the gastrointestinal tract is essential for gut health, providing a protective barrier against pathogens while maintaining symbiosis with the microbiome. Its disruption is a hallmark of gastrointestinal diseases like ulcerative colitis. While glycan foraging by gut bacteria is thought to initiate mucus disruption, its impact on mucus structural properties remains poorly understood, largely due to the lack of physiologically relevant models. To address this gap, we developed a method to collect human-cell-derived mucus that closely mimics the mechanical properties of human colonic mucus. Using this system, we investigated mucus utilization and degradation by a panel of commensal bacteria with distinct metabolic profiles. Glycan utilization by species such as *Bacteroides thetaiotaomicron* and *Bacteroides fragilis* showed no correlation with changes in mucus rheology. Instead, secreted proteases were identified as the primary driver of mucus degradation. Protease activity by *B. fragilis* and *Bifidobacterium longum* was influenced by nutrient availability, whereas in *Enterococcus faecalis,* it was additionally affected by oxygen exposure*. E. faecalis* also adapted to oxidative stress by enhancing carbohydrate metabolism and upregulating several virulence genes. Together, our findings reveal that bacterial mucus degradation is context-dependent and shaped by environmental factors. This study provides key insights into the mechanisms underlying mucus degradation and underscores the value of human cell-derived mucus models for understanding bacteria-mucus interactions in health and disease.

## INTRODUCTION

The mucus layer lining the gastrointestinal tract protects the gut by lubricating and shielding the epithelium from abrasion, pathogens, and harmful substances, while aiding nutrient absorption^1^. This layer also supports a balanced relationship between the microbiome and the host, with commensal bacteria residing in the top portion of the mucus and influencing mucus production^2^, immune system development^3^, and pathogen defense^4^. Animal models demonstrate the importance of the mucus layer in gut health, as deficiencies in mucus secretion cause colitis and increase infection susceptibility^5^. As such, disrupted mucus integrity is a hallmark in patients with ulcerative colitis and Crohn’s disease^6,7^. In ulcerative colitis, the mucus layer is often thinner in inflamed areas, accompanied by an increase in mucosa-associated bacteria and even bacterial penetration into the epithelium^8^.

The function of intestinal mucus relies on its viscoelastic properties, primarily governed by mucins^9^. Mucins are large extracellular glycoproteins, consisting of approximately 80% carbohydrates in a bottle-brush configuration around a protein core, with globular hydrophobic regions^9^. The N- and C-terminal cysteine-rich domains within the hydrophobic regions facilitate mucin polymerization through intermolecular disulfide bonds and hydrophobic interactions^9^. These interactions regulate the density of crosslinks, mucus swelling, mesh size, and transport characteristics^9,10^.

Commensal gut bacteria encode glycosyl hydrolases with broad glycan-degrading capabilities, allowing them to utilize nearly all major plant and host glycans, including those in mucus^11,12^. This glycan foraging by commensal bacteria is considered an initial stage in mucus disruption^11,13^. However, the impact of mucin glycan utilization on mucus viscoelastic properties and whether glycan foraging alone causes mucus degradation is unknown. Rather, other mechanisms may be at play. For example, *Entamoeba histolytica,* the parasite responsible for amoebiasis, secretes a cysteine protease that disrupts the mucus structure via proteolytic cleavage, enabling parasite invasion^14^. Alternatively, *Helicobacter pylori* reduce mucus viscosity by increasing pH, which alters gastric mucus rheology and enhances bacterial motility for colonization^15^.

Ideally, one would use intact human mucus to examine changes in mucus structural properties. However, obtaining sufficient native human colonic mucus is challenging. Gnotobiotic animals offer advantages for studying bacteria-mucus interactions, but the complexity of the mucus microenvironment makes isolating specific interactions *in vivo* challenging^16^. As a result, researchers often use porcine gastric mucin (PGM) as a cost-effective, commercially available alternative. Although PGM is the current gold standard for studying bacteria-mucus interactions, it fails to accurately replicate the structural and mechanical properties of native human colonic mucus, which exists at near-neutral pH. Even when extracted with the least degradative purification protocols performed by individual laboratories^17^, reconstituted PGM from these non-commercial sources behaves as a gel only at low pH (<4)^18^, while commercial PGM fails to gel at any pH^19^. This limits its suitability for studying gut commensal bacteria-mucus interactions under physiological conditions. Hybrid systems, such as mucin-based hydrogels that combine purified mucins with components like alginate^20^ or methylcellulose^21^, and mucin-inspired hydrogels based on synthetic materials like dendritic polyglycerol sulfate^22^, are similarly suboptimal. Their properties are largely dominated by non-mucin materials, resulting in significant deviations from native mucus^23^. Additionally, mucin-inspired hydrogels lack the oligosaccharide components essential for understanding how glycan utilization impacts mucus structural properties^22^.

To overcome these limitations, we optimized a method for harvesting human cell-derived mucus (HCDM) and found that it recapitulates human colonic mucus. Unlike PGM, HCDM forms a soft, three-dimensional (3D) viscoelastic gel at physiological pH, and therefore serves as a more biological model for native mucus in studies of bacteria-mucus interactions. Leveraging the HCDM platform, we investigated how gut commensal bacteria influence the viscoelastic properties of mucus. Our results reveal that while glycan utilization alone does not alter the mechanical integrity of mucus, bacterial proteases play a pivotal role in disrupting its network. This protease activity is regulated by environmental factors such as nutrient availability, particularly by *Bacteroides fragilis* and *Bifidobacterium longum*, and nutrient availability and oxygen levels by *Enterococcus faecalis*. In-depth analysis of *E. faecalis*, a pathobiont, uncovered transcriptional shifts in virulence factors that enable it to adapt and degrade mucus under specific conditions, highlighting its survival strategies within the host. These findings provide valuable insights into the intricate interactions between the gut microbiome and the mucus barrier, emphasizing the necessity of using physiologically relevant mucus models for such studies.

## RESULTS

### Human cell-derived mucus (HCDM) mimics human colonic mucus

Goblet cells are specialized epithelial cells that reside within the intestinal epithelium. They continuously produce mucus, which is stored in granules within their cytoplasm and then excreted after mucus-filled granules fuse with the cell membrane^24^. This process is accelerated by cholinergic agonists, which induce mucus discharge by activating adenylyl cyclase receptors or voltage-dependent calcium channels^25,26^. By employing different cholinergic agonists, we were able to harvest mucus secreted by HT29-MTXE12 goblet cells as a gel and compare its mechanical properties to those of human colonic mucus.

To maximize the mucus harvest, we tested four cholinergic agonists: forskolin, 3-isobutyl-1-methylxanthine (IBMX), carbachol, and bethanechol. Forskolin and IBMX stimulate exocytosis through a cAMP-dependent mechanism, while carbachol and bethanechol work by increasing cytosolic calcium levels, which induce mucus discharge^24,25^. Individual secretagogues were added to HT29-MTXE12 cells after three weeks of culture, at the peak of mucus accumulation *in vitro* reported in the literature^27^ and observed experimentally (**Fig. 1A**). All treatments facilitated mucus removal without disturbing the underlying cell monolayer (**Fig. 1B, C**). However, mucus harvested using forskolin delaminated more easily and cohesively with gravity (**Supplementary Fig. S1A**), while the other secretagogues produced a more fragmented delamination. Forskolin also yielded the highest dry mass per surface area compared to the other secretagogues (**Fig. 1D).**

**Figure 1.**
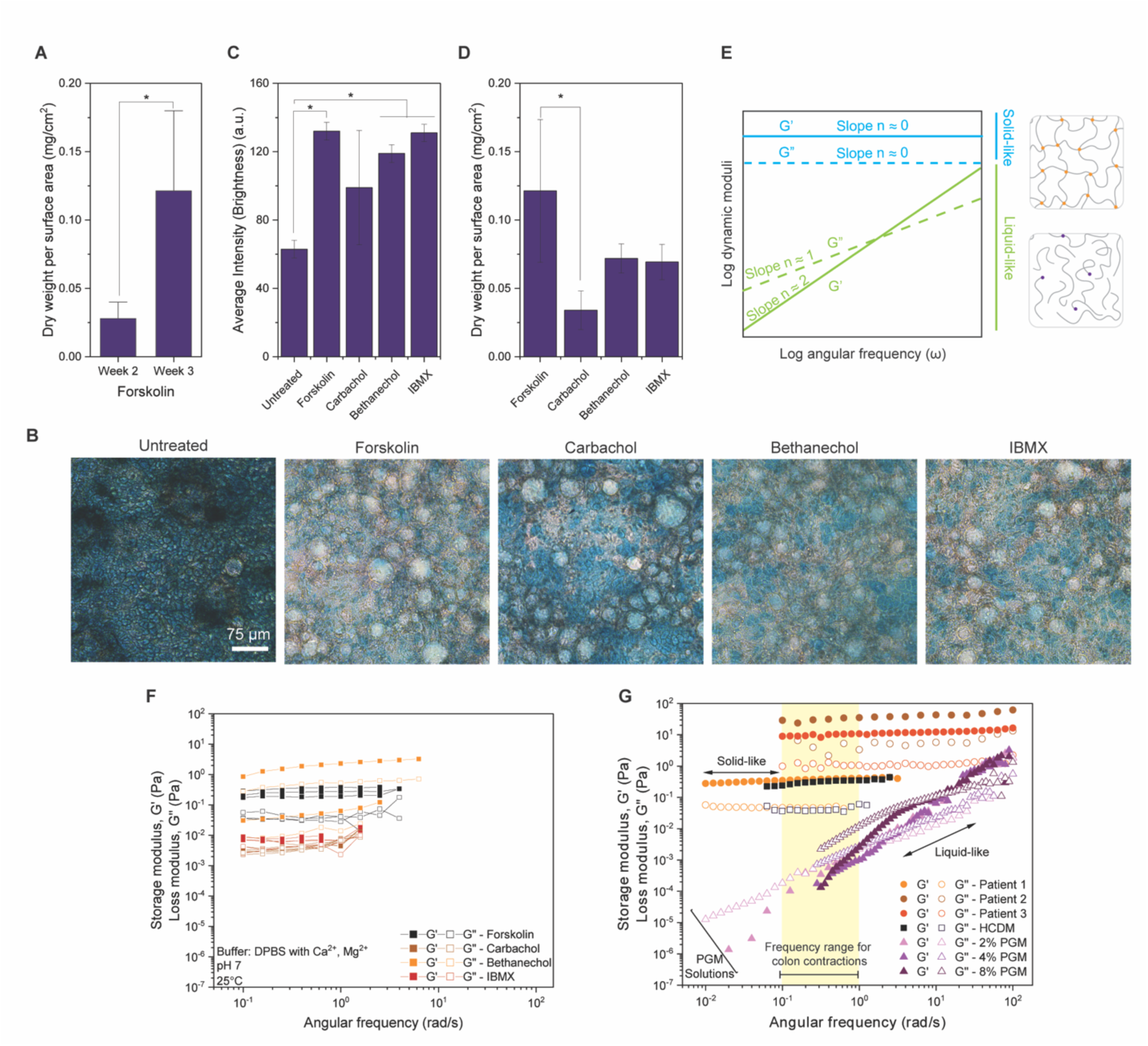
Human cell-derived mucus (HCDM) as a model for human colonic mucus. **(A)** Mucus yield (dry mass per unit area) measured after two and three weeks in culture. **(B)** Micrographs of HT29-MTX-E12 goblet cells stained with Periodic acid-Schiff (PAS)/Alcian blue before and after treatment with forskolin, carbachol, bethanechol, or IBMX, showing undisturbed cell monolayers following mucus exocytosis. **(C)** Average staining intensity (brightness) of residual mucus post-collection compared to the untreated control. **(D)** Average dry mass per unit area of harvested HCDM for each secretagogue. **(E)** Characteristic curves for oscillatory shear rheology, illustrating solid-like elastic gels and liquid-like viscous solutions, with expected slope values at low frequencies on a logarithmic plot. **(F)** Oscillatory shear rheology performed on HCDM harvested using different secretagogues. **(G)** Mechanical spectrum of HCDM (collected with forskolin) compared to human colonic biopsies and PGM solutions at different concentrations. The yellow region in the plot indicates the frequency range for mixing and segmentation contractions in the colon^36^. All bars represent mean ± SD. Asterisks denote statistical significance: *p* < 0.05 (*).

Next, we used oscillatory shear rheology to evaluate the impact of each secretagogue on the mucus mechanical properties. Oscillatory rheology measures the viscous and elastic properties (dynamic moduli) of a material, characterized by the storage (G’) and loss (G”) modulus at various frequencies^28^. In a typical rheological experiment, the material is subjected to cyclic mechanical deformation, with G’ indicating resistance to elastic deformation and G” reflecting resistance to flow (**Fig. 1E**). This information enables discrimination between gels (formed by crosslinked networks) and viscous solutions, based on the slopes and magnitudes of G’ and G”, as well as the ratio of viscous to elastic effects (phase angle, (G”/G’)). Elastic gels exhibit constant G’ and G” values regardless of frequency, with G’ much larger than G”. By contrast, viscous solutions have higher G” than G’ at low frequencies and show power-law behavior^28^, with G’ increasing twice as fast as G” (**Fig. 1E)**. Forskolin yielded the greatest consistency in mechanical properties across batches, with an average standard deviation of 0.072, compared to 1.31 for bethanechol, which also produced elastic gels (**Fig. 1F**). Given the increased yield and mechanical reproducibility of the mucus collected with forskolin, we used this secretagogue in all subsequent experiments.

HCDM closely matched the mechanical spectrum of mucus obtained from three colorectal cancer patients’ colonic resections, particularly overlapping with mucus from patient 1 (**Fig. 1G**). Direct comparisons to healthy colonic mucus remain challenging due to the unavailability of resections from healthy individuals. However, previous studies have shown that MUC2/MUC5 expression ratios in HT29-MTX-E12 cells vary depending on culture conditions^29,30^. MUC2 is the major structural component of healthy colonic mucus, whereas MUC5 is associated with inflammatory states like IBD^31^. In our earlier work, we found that under our experimental conditions, HT29-MTX-E12 cells exhibited higher MUC2 expression compared to MUC5AC and MUC5B mucins^32^. LS174T cells, an alternative colorectal adenocarcinoma cell line, which predominantly express MUC2^33^, and differentiated human primary colonic epithelial cells failed to secrete sufficient mucus for analysis, even after treatment with DAPT and Y-27632 to induce secretory cell marker expression^34^.

Unlike reconstituted extracellular matrices like type-I collagen, basement membranes and fibrin, whose G’ and G” oscillate between 5 to 100 Pa^35^, HCDM is extraordinarily compliant, with an average G’ of 0.57 ± 0.47 Pa and G” of 0.09 ± 0.08 Pa at 1 rad/s and pH 7.0. Rheology of both HCDM and human biopsies resulted in curves characteristic of elastic gels, and values for the phase angle close to zero (**Supplementary Fig. S1B**). Conversely, commercial PGM did not replicate human colon mucus’s mechanical properties, showing liquid-like behavior typical of viscous solutions (**Fig. 1G)**.

### Mechanical properties of HCDM are robust to changes in pH

Mucus naturally experiences significant pH variation along the digestive tract in healthy individuals: it is highly acidic in the stomach (1.0-2.5), increases to 6.6 in the proximal small intestine, rises to 7.5 in the terminal ileum, drops sharply to 6.4 in the caecum, and then progressively increases from the right ascending colon (6.4) to the left descending colon (7.0)^37^. These pH changes strongly affect PGM’s rheology, causing sol-to-gel transitions, from a solution at pH 6 to a gel at pHs 2 and 4^10,18^. The gelation of PGM under acidic conditions is driven by reduced electrostatic repulsion between highly negatively charged glycan segments, such as those containing sialic acid, thereby increasing hydrophobic aggregation within the macromolecule^38^. Since the pH of the culture media also fluctuates during bacterial growth *in vitro* due to sugar fermentation, we investigated HCDM’s rheology at acidic, neutral, and alkaline conditions.

HCDM proved highly resistant to changes in pH (**Fig. 2A**). Across a pH range of 2–6, its dynamic moduli showed no sol-to-gel transitions and a notable increase in the storage modulus, contrasting the dramatic 1000-fold shift reported for reconstituted PGM at the same frequency^18^. This resilience likely stems from limited chain rearrangement in native mucus compared to reconstituted PGM^18^. At pH 2, the storage modulus increased 2.4-fold compared to pH 6, with no significant changes in the loss modulus (**Fig. 2B**). These results are consistent with findings for cervicovaginal mucus, which shows minimal pH-dependent changes^38^. Notably, there were no significant differences in the dynamic modulus between pH 7 and pH 9, suggesting that disulfide crosslinks maintain a stable elastic network, which is reinforced by non-chemical crosslinking interactions at lower pH^38^. We also found that the mechanical properties of HCDM remained unchanged after being frozen at -20/80°C for 1 week and then thawed to room temperature (**Supplementary Fig. S2)**.

**Figure 2.**
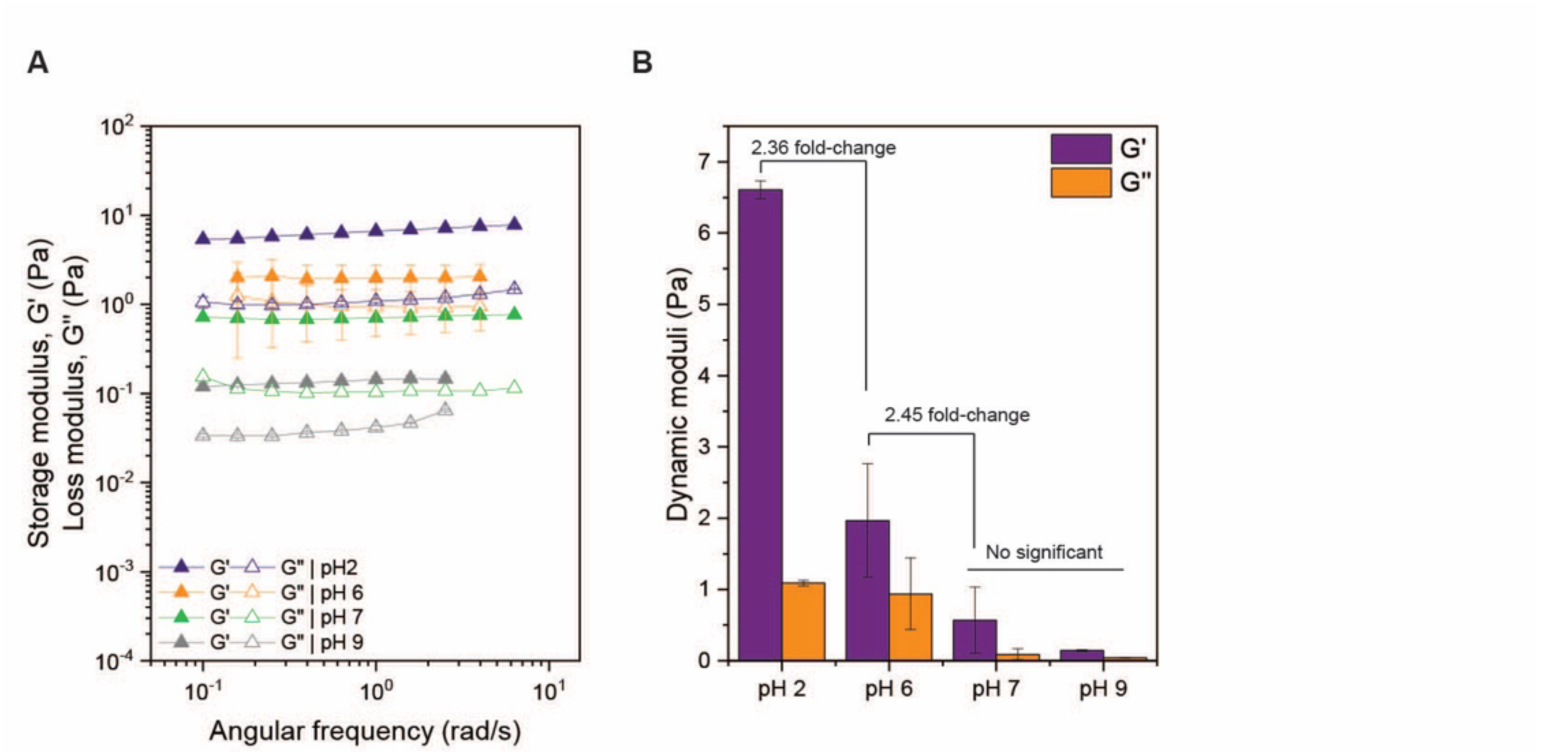
HCDM is exceptionally soft, yet its gel-like elasticity is robust to pH variations. **(A)** Small-amplitude oscillatory rheology was performed across a pH range of 2 to 9. **(B)** Storage modulus (G’) and loss modulus (G”) values at 1 rad/s are shown for each pH condition.

### Glycan utilization does not alter mucus viscoelasticity, whereas proteases do

To investigate how commensal gut bacteria influence mucus mechanical properties, we selected a representative panel of gut commensals with distinct metabolic profiles (**Fig. 3A**). This panel included bacteria capable of digesting complex carbohydrates, including mucins, such as *Bacteroides thetaiotaomicron*, *Bacteroides caccae*, and *Bacteroides fragilis*^39,40^. Other members, including the butyrate producers *Faecalibacterium duncaniae*, *Roseburia intestinalis*, and *Ruminococcus gnavus*, specialize in targeting specific components of the mucus, such as sialic acid^41^ and cross-feeding on mucin O-glycans^42,43^. Additionally, we included *Enterococcus faecalis*, a pathobiont linked to systemic infections^44^, and *Bifidobacterium longum subsp. Infantis* (*B. longum*), a probiotic known for foraging on mammalian oligosaccharides^45^, to capture a range of mucin and oligosaccharide-degrading capabilities.

**Figure 3.**
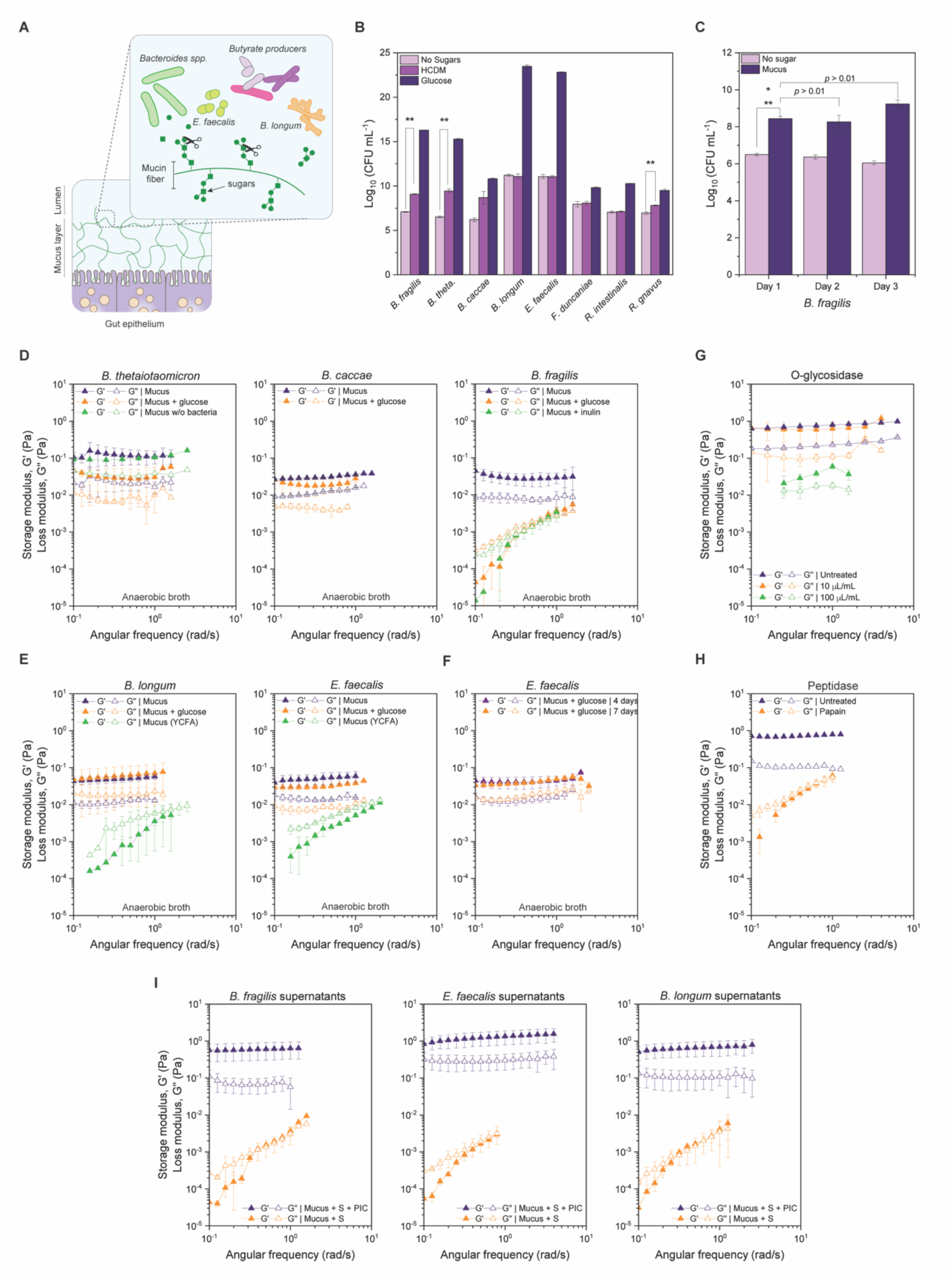
Influence of commensal gut bacteria on mucus utilization and structural integrity. **(A)** Commensal gut bacteria with distinct metabolic profiles were cultured with HCDM as the sole carbon source or in combination with alternative carbon sources for 24 hours to evaluate their impact on mucus structural properties. **(B)** Bacterial growth was quantified by counting colony forming units (CFUs) after 24 hours in HCDM or glucose. **(C)** Growth of *B. fragilis* was measured by CFU counts over 1, 2, and 3 days in HCDM. **(D, E)** Bulk rheological properties of mucus were analyzed after co-culture with mucolytic bacteria in HCDM supplemented with glucose or inulin. Mucus incubated without bacteria under the same conditions served as a control. The drop in dynamic moduli is caused by reducing agents in the anaerobic broth (i.e., cysteine), required for cultivating strict anaerobes. **(F)** Rheological analysis of *E. faecalis* performed after incubation with mucus supplemented with glucose after 4 or 7 days. **(G, H)** Oscillatory rheology on mucus treated with glycolytic enzymes (O-glycosidase) or the proteolytic enzyme papain **(I)** Mechanical spectrum of HCDM treated with bacterial supernatants from *B. fragilis, B. longum,* and *E. faecalis,* with and without a protease inhibitor cocktail. PIC: protease inhibitor cocktail, S: cell-free supernatants. All bars represent mean ± SD. Asterisks denote statistical significance: *p* < 0.05 (*), *p* < 0.01 (**).

Bacteria were cultured in chemically defined broths with mucus as the sole carbon source for 24 hours, except for *F. duncaniae*, *R. intestinalis*, and *R. gnavus*, which were cultured in glucose-free yeast-casitone-fatty-acids (YCFA) broth. We used agar plating to quantify mucus utilization via colony forming units (CFU) per mL and small-amplitude oscillatory rheology to assess changes in mucus structural properties. We also included mucus samples without bacteria as controls in all experiments to effectively distinguish media-induced changes from bacterial degradation.

Among the tested species, *B. thetaiotaomicron* and *B. fragilis* showed significant but modest growth on mucus, substantially lower than on glucose (**Fig. 3B**), in agreement with previous studies^46^. Extended incubation of *B. fragilis* for 3 days did not increase CFU counts (**Fig. 3C**), suggesting limited glycan utilization. Neither *B. longum* nor *E. faecalis* utilized mucus, despite possessing pathways associated with mucin foraging, such as the Family 1 of solute binding proteins and extracellular sulfatase domains in *B. longum*^45^ and *E. faecalis* ^47^, respectively.

None of the bacteria, including the mucolytic species like *B. thetaiotaomicron*, *B. caccae*, and *B. fragilis*, altered mucus rheological properties after 24 hours (**Fig. 3 D, E; Supplementary Fig. S3**). Given that dietary factors influence glycan foraging^48^, we hypothesized that the presence of alternative carbon sources may affect the ability of these organisms to utilize mucus. We therefore supplemented the chemically defined broth and YCFA with glucose and reassessed changes in mucus rheological properties. At 24 h, the mucus retained its gel-like properties in all cases except for *B. fragilis* (**Fig. 3D, E**; **Supplementary Fig. S3**), which solubilized it into a liquid-like solution. Substituting glucose with inulin, a metabolizable polysaccharide for *B. fragilis*, produced similar degradation (**Fig. 3D**). This behavior persisted in YCFA and PYG broths (**Supplementary Fig. S4**), indicating that *B. fragilis* enhances mucus degradation in nutrient-rich conditions with both simple and complex sugars.

Conversely, mucus degradation by *B. longum* and *E. faecalis* was suppressed in glucose-containing broths (**Fig. 3E)**. This suppression was not due to slow degradation, as mucus remained intact after 7 days of incubation with *E. faecalis* (**Fig. 3F**). Similarly, replacing glucose with lactose or galactose also inhibited mucus degradation by *E. faecalis* (**Supplementary Fig. S5**). However, mucus degradation occurred in YCFA without glucose, an undefined media containing proteolytically degradable nitrogen sources, among other nutrients (**Fig. 3E**). In contrast, *F. duncaniae, R. intestinalis*, and *R. gnavus* did not degrade mucus under any tested condition (**Supplementary Fig. S3**), with only *R. gnavus* showing slight utilization (**Fig. 3A**).

Enzymatic removal of mucus glycans with O-glycosidases, sialidases, and fucosidases did not significantly affect mucus bulk rheological properties (**Fig. 3G, Supplementary Fig. S6**). O-glycosidase cleaves core 1 and core 3 O-linked glycans, which are built on GalNAc residues and extended by sugars into linear or branched chains, with core 1 prevalent in mucins secreted by HT29-MTX-E12 cells^49^. Sialidase and fucosidase target terminal sialic acid residues and α1-3/α1-4 linked fucose residues in mucins, respectively. Even at high enzyme concentrations, although the dynamic moduli decreased, the mucus gel-like structure was retained (**Fig. 3G, Supplementary Fig. S6**). In contrast, treatment with papain, a cysteine protease with broad specificity, solubilized the mucus, as evidenced by frequency-dependent changes in G’ and G” (**Fig. 3H**).

We therefore hypothesized that mucus solubilization by *B. fragilis*, *B. longum*, and *E. faecalis* resulted from proteolytic cleavage. Previous studies identified papain-like cysteine proteases in *Bacteroides* species, including *B. fragilis*^50^, whose activity increases in nutrient rich conditions^51^. Similarly, *B. longum* encodes endopeptidases with homology to enzymes in *Bifidobacterium animalis* capable of casein hydrolysis^52^, a pre-digested form of casitone present in YCFA. *E. faecalis*, on the other hand, produces metalloproteinases like gelatinase, which are known to degrade collagen^53^.

To test this hypothesis, we measured protease activity in whole cells and cell-free supernatants using azocasein to determine their localization. In *B. fragilis*, protease activity was detected in both whole cells and cell-free supernatants in cells grown in defined minimal media with glucose, consistent with previous findings^51^ (**Supplementary Fig. S7**). In contrast, *B. longum* and *E. faecalis* exhibited significantly higher protease activity in cell-free supernatants compared to whole cells when cultured in glucose-free YCFA (**Supplementary Fig. S7**). Importantly, treating mucus with cell-free supernatants from *B. fragilis*, *B. longum*, and *E. faecalis* caused degradation, which was inhibited by a protease inhibitor cocktail **(Fig. 3I).** These results confirm that these species disrupt intestinal mucus through proteolytic enzyme secretion, rather than glycan foraging.

### *E. faecalis* upregulates mucus degradation genes in the presence of oxygen and glucose

Given the capacity of *E. faecalis* to cause extraintestinal infections, we hypothesized that it may employ mucolytic traits in the presence of oxygen. Oxygenation is known to enhance colonization by other opportunistic pathogens within the host. For instance, *S. pneumoniae* binds more effectively to epithelial cells under aerobic conditions^54^, and increased oxygenation near the GI mucosa enhances the virulence of *Shigella flexneri*, promoting epithelial cell invasion^55^. While several virulence factors of *E. faecalis*—including surface adhesins, gelatinase, superoxide production, cytolysin, and hemolysins—have been identified^56,57^, the environmental factors regulating these traits are not well understood. Indeed, oxygen exposure triggered mucus degradation by *E. faecalis* regardless of glucose supplementation, unlike the intact mucus observed under anaerobic conditions **(Fig. 4A).**

**Figure 4.**
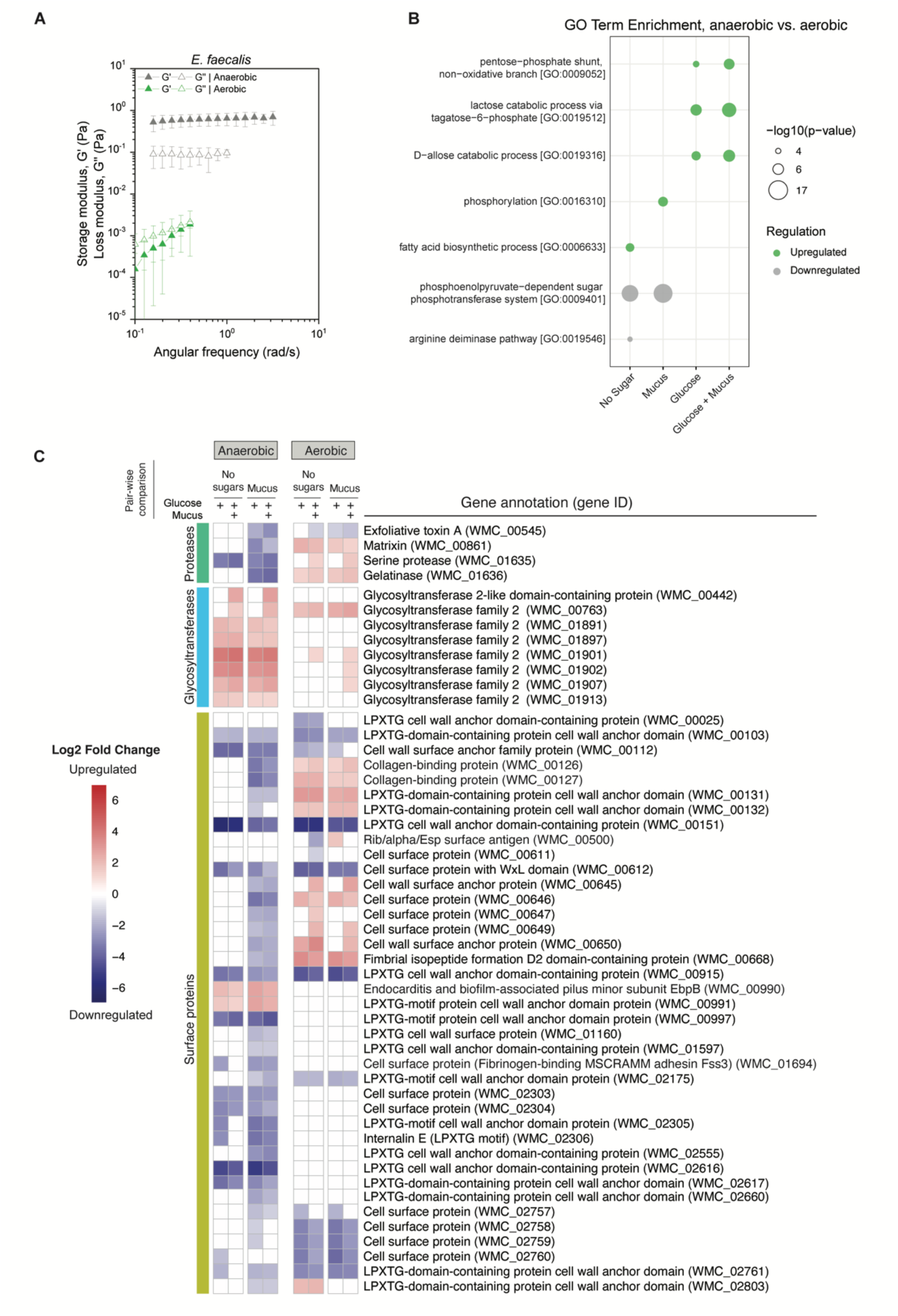
Impact of oxygen and glucose on *E. faecalis*-mediated mucus degradation and gene expression. **(A)** Transcriptomics was performed on *E. faecalis* grown in anaerobic or oxygenated conditions with or without mucus. Differential gene expression analysis was conducted using DESeq2 with an adjusted *p-value* threshold of <0.05 and a |log₂FoldChange| > 1. **(B)** Pathway analysis using GO term enrichment was performed on *E. faecalis* cells grown in oxygenated versus anaerobic conditions. **(C)** Genes involved in virulence were selected for inclusion in the heatmap, which displays their relative expression levels across conditions.

To investigate the mechanisms driving *E. faecalis*–induced mucus degradation, we examined the effects of oxygen and sugar availability on gene expression. Under aerobic conditions with glucose, genes involved in carbohydrate metabolism, including lactose and D-allose catabolic processes, and the non-oxidative pentose-phosphate pathway (PPP), were upregulated (**Fig. 4 B**). These pathways support energy production via ATP generation and detoxification of reactive oxygen species through glutathione synthesis^58,59^. Conversely, in glucose-starved cells, *E. faecalis* responded to oxygen by upregulating fatty acid biosynthesis genes, while downregulating the phosphoenolpyruvate (PEP)-dependent sugar phosphotransferase system (PTS) and the arginine deiminase pathway, indicating a metabolic shift towards more efficient aerobic respiration (**Fig. 4 B**). This is consistent with previous studies showing that oxidative stress and glucose limitation induce membrane remodeling through increased unsaturated fatty acids, which offer better antioxidant protection^59^. When mucus served as the primary carbon source, phosphorylation but not fatty acid biosynthesis was upregulated, suggesting the activation of oxidative phosphorylation and stress response mechanisms^59^ (**Fig. 4B**).

Oxygen and glucose also triggered the expression of virulence genes, including proteases like matrixin, gelatinase, and serine proteases (**Fig. 4C, Supplementary Fig. S8A)**. These enzymes are key to enterococcal-mediated diseases due to their tissue-destructive capabilities and interference with host-cell signaling^60,61^. In particular, gelatinase and matrix metalloproteinases (MMPs), or matrixins, are implicated in the pathogenesis of inflammatory bowel disease (IBD)^62,63^. This is supported by increased expression in oxygenated conditions, in which mucus is degraded, compared to anaerobic conditions, during which the mucus remains intact (**Fig. 4A**). Oxygen and glucose also influenced surface protein expression, upregulating collagen-binding proteins (**Fig. 4C**), probably to enhance adhesion in oxygen-rich environments, such as those outside the gastrointestinal tract.

In anaerobic, glucose-rich conditions, in which mucus is not degraded, *E. faecalis* significantly upregulated glycosyltransferases **(Fig 4C**). These enzymes are involved in the biosynthesis of structural polysaccharides and glycoconjugates^64^ as well as the formation of cell walls, capsules, and other extracellular polysaccharides that facilitate bacterial adhesion and colonization^65,66^. When both glucose and mucus were present, two specific glycosyltransferases were upregulated. One gene shared 90% homology with a hyaluronan synthase involved in glycosaminoglycan synthesis, a capsular polysaccharide that enhances the infectivity of Group A *Streptococci* by resisting neutrophil-mediated killing^67^. The other showed homology to a glycosyltransferase linked to biofilm formation in *Lactobacillus divergens* and *Paenibacillus alvei*.

Notably, fewer surface proteins were induced in anaerobic, glucose-rich conditions, with notable exceptions like the endocarditis and biofilm-associated pilus (EbpB) (**Fig. 4C**). Ebp structures promote cell adhesion and are considered key triggers for biofilm formation in *E. faecalis*^68^. Despite this, minimal biofilm formation was observed via crystal violet staining (**Supplementary Fig. S9A**). In contrast, lectin staining with concanavalin A revealed exopolysaccharides that colocalized with bacterial cells (**Supplementary Fig. S9 B**). These findings suggest that these glycosyltransferases may facilitate microcolony formation rather than extensive biofilm development. Interestingly, alternative sugar pathways suppressed by glucose alone were reactivated when mucus was added (**Supplementary Fig. S8B**), potentially reflecting increased sugar demand.

In anaerobic conditions with mucus as the primary carbon source, *E. faecalis* exhibited a distinct expression profile characterized by the upregulation of LPxTG-motif cell wall-anchored proteins, including collagen- and fibrinogen-binding proteins (**Fig. 4C**). These surface proteins are critical in enterococcal infections by mediating adherence to host tissues, a key step for colonization and infection^69^. Proteases, such as matrixin, gelatinase, and a serine protease homologous to *S. aureus* exfoliative toxin A, were also induced in these conditions, coinciding with the mucus degradation observed previously under these conditions (**Fig. 3E**). Notably, hemolysin—a cytolytic enzyme^57^—was upregulated (**Supplementary Fig. S8A**), further linking oxygen availability to virulence expression.

## DISCUSSION

Accurate mucus models that capture the physical properties of mucus are essential for understanding how the gut microbiome impacts mucus integrity. Here, we used human cell-derived mucus or HCDM, a model that closely mimics the mechanical properties of human colonic mucus. HCDM is exceptionally soft, mechanically consistent, sterile, and free from microbial and food contaminants, making it a reliable tool for investigating bacteria-mucus interactions. Additionally, HCDM demonstrates greater tolerance to acidification compared to reconstituted PGM^18^. Previously, we demonstrated the utility of HCDM by investigating the diffusion of non-mucoadhesive nanoparticles^70^ and microplastics^32^, showing that its mesh size (214.04 ± 4.29 nm at pH 7) effectively immobilizes nanoparticles of 500 nm in diameter. This finding agrees with reported pore size distributions in pig jejunum mucus (20-200 nm)^71^ and human colonic mucus (below 500 nm)^72^, further validating HCDM as an accurate model for mucus studies.

This study addresses a key question: does bacterial metabolism of mucus alter its structural properties? While some bacterial species are known to utilize mucin-decorating sugars^73^—a process traditionally seen as a primary driver of mucus degradation^12^ —our findings indicate that such glycan utilization does not necessarily disrupt the mucus network. For example, mucolytic species such as *B. thetaiotaomicron* and *B. fragilis* exhibited significant growth after 24 hours on mucus. Similarly, butyrate-producing bacteria and species with oligosaccharide-degrading capabilities, such as *B. longum* and *E. faecalis*, showed neither significant mucus utilization nor alterations to its rheological properties under similar conditions. These results align with previous studies showing that bacterial growth preferences vary depending on mucin source^74^, with colonic mucus being more resistant to utilization^75^. However, growth on mucus did not always result in changing its rheological properties, as illustrated by *B. thetaiotaomicron* and *B. fragilis*.

Enzymatic removal of mucin O-glycans also failed to impact HCDM’s bulk rheology, consistent with prior studies showing minimal effects of terminal sialic acid removal on bovine cervical mucus properties^76^. These findings underscore the resilience of colonic mucus despite its utilization by bacteria. However, papain treatment led to HCDM solubilization, paralleling earlier reports of papain-induced solubilization of intact human gastric mucus under analogous experimental conditions^77^. Interestingly, supplementation with either glucose or inulin triggered mucus degradation by *B. fragilis*, while degradation by *B. longum* and *E. faecalis* occurred in YCFA, an undefined media containing proteolytically degradable nitrogen sources.

The loss of mucus degradation in the presence of protease inhibitors further supports the idea that proteases, rather than glycoside hydrolases, are responsible for compromising mucus structure. Although bacterial proteases are generally associated with enhancing nutrient acquisition through proteolysis, they have been also shown to contribute to host tissue damage during infections by degrading connective tissues and facilitating bacterial spread^53,63,78,79^. It is also plausible that glycan utilization by commensal bacteria increases the exposure of mucus to proteases, thereby weakening its barrier properties.

The mucolytic activity of *B. fragilis* is particularly notable when compared to other *Bacteroides* species (i.e., *B. thetaiotaomicron* and *B. caccae*). Like closely related species, *B. fragilis* is widely recognized for its polysaccharide-degrading activities, which play a key role in nutrient acquisition within the gut^46^. However, its proteases have been linked to fibrinogen hydrolysis^80^ and implicated in diarrheal disease^81^. Notably, *B. fragilis* produces fragipain, a protease essential for activating *B. fragilis* toxin (BFT), which is associated with colitis and intestinal malignancies^82^. These proteases have also been detected in outer membrane vesicles^83^, although their ecological function within the gut remains poorly understood.

In *E. faecalis*, oxygen exposure also triggered mucus degradation, independent of glucose supplementation, and coincided with the upregulation of gelatinase, matrixin, and serine proteases. In anaerobic conditions, when mucus was the primary carbon source, *E. faecalis* upregulated multiple surface proteins, including collagen- and fibrinogen-binding proteins, consistent with previous studies linking surface protein expression to stressors like glucose starvation^84^, sub-lethal heat stress^85^, and the presence of extracellular matrix molecules^86^. When both glucose and mucus were present in anaerobic conditions, glycosyltransferase expression increased, accompanied by biofilm-associated pilus upregulation and increased sugar metabolism. These adaptations coincided with the preservation of mucus integrity, probably linked to exopolysaccharide production and microcolony formation. Such metabolic flexibility and virulence regulation likely explain *E. faecalis* capacity to efficiently colonize mucosal environments where nutrient sources fluctuate, and oxygen concentrations vary. Specifically, the increase in energy yield under oxygen may explain why restricting sugar intake limits *E. faecalis* growth in antibiotic-treated mice^87,88^. It also sheds light on the predominance of *E. faecalis* at colon reconnection sites, where collagen-degrading activity of *E. faecalis* is elevated^53^.

While this study provides valuable insights into bacteria-mucus interactions, we have identified some limitations. First, albeit oscillatory rheology is highly sensitive to structural changes in soft polymers, it may not capture local variations in the mechanical properties of mucus. This challenge is further compounded by the composition of anaerobic broths used to cultivate fastidious bacteria, which may introduce artifacts unrelated to bacterial activity. To mitigate this, we used control mucus samples without bacteria and focused on events leading to total network disruption (gel-to-liquid transition). Second, HCDM contains a mixture of primarily MUC5AC, MUC5B, and MUC2^32^, the latter being dominant in healthy colonic mucus. The impact of these compositional differences on mucus degradation is unclear. Despite these challenges, our findings align with previous research showing the susceptibility of human gastric mucus to proteolytic degradation^77^, and the resistance of colonic mucus to bacterial utilization compared to PGM^74,75^. Lastly, our study does not consider the role of other mucosal components^89^, such as antimicrobial peptides, immunoglobulins, trefoil factors, and the interactions among microorganisms within mucus.

In summary, we have optimized a method for collecting human-cell-derived mucus that closely mimics the mechanical properties of human colonic mucus. This mucus is exceptionally soft yet is robust to changes in pH. Using this system, we investigated mucus utilization and degradation by a panel of gut commensals with distinct metabolic profiles. Our findings revealed that bacterial metabolism of mucus is not always tied to its structural disruption, with many commensals, including *B. fragilis* and *E. faecalis*, exhibiting specific nutrient- and oxygen-dependent protease activity as key drivers of degradation. Protease inhibitors effectively prevented mucus breakdown, underscoring the role of proteolytic enzymes in mucus disruption. Additionally, *E. faecalis* demonstrated metabolic flexibility and virulence factor regulation in response to fluctuating environmental conditions, including oxygen and nutrient availability. While offering new insights into bacteria-mucus interactions, this study acknowledges limitations, including the limited representation of MUC2 in HCDM. However, our findings provide a foundation for exploring how environmental and microbial factors influence the mucosal barrier, with implications for understanding gut health and disease.

## MATERIALS AND METHODS

### Mammalian cell culture and mucus collection

HT29-MTX-E12 cells were routinely cultured on T-25 flasks with Dulbecco’s modified Eagle’s medium (DMEM) supplemented with 10% fetal bovine serum. The medium was changed every other day for 21 days. To collect intact mucus, we modified the method used by Capon and coworkers^71^. On day 21, the flasks were rinsed with Dulbecco’s phosphate-buffered saline (DPBS) without calcium and magnesium to remove culture media, and incubated with 1 mM of forskolin, carbachol, bethanechol, or 3-isobutyl-1-methyl-xanthine (IBMX) for 1 h. The cells were then gently rinsed up to five times with DPBS. After that, 1 mL of fresh DPBS (with calcium and magnesium) or broth was used to gently flush the cell monolayer and delaminate the mucus. To determine total mass based on dry weight, cells were rinsed with deionized water instead of DPBS several times, and the mucus was collected in the same water, frozen overnight at –20°C, lyophilized, and weighed using a microbalance. All treatments were performed in triplicates.

### Periodic acid-Schiff (PAS)/Alcian blue staining

We employed a PAS/Alcian Blue stain kit (Epredia) to detect both neutral and acidic mucus secretions before and after secretagogue treatment. Briefly, flasks containing 21-day-old HT29-MTX-E12 cells, with mucus (untreated) or without mucus (post-secretagogue exposure), were fixed with 10% formalin for 2 h. After fixation, the flasks were washed several times with deionized water and stained with Alcian Blue for 15 minutes, followed by rinsing with deionized water. Subsequently, the cells were treated with periodic acid solution for 5 minutes, stained with Schiff’s reagent for 15 minutes, and then rinsed again. The stained cells were immediately imaged using a brightfield microscope at 20X magnification.

### Human colonic mucus

Human participant research was approved by Cornell University’s Institutional Review Board (IRB) under the protocol number 19-04020223. All participants provided informed consent prior to their operations. Human colonic biopsies were obtained from surgical resections of cancerous tissue. Mucosal samples were collected by scraping the luminal contents from the apical surface of the resected tissue. These samples were placed in sterile cryovials, stored on dry ice, and transported to the laboratory. Upon arrival, they were stored at - 80°C until use. Just before mechanical characterization, the resected tissue, including HCDM samples, was equilibrated in DPBS with calcium and magnesium (pH 7) for 1 h at room temperature using a 100 kDa dialysis back and then characterized by dynamic oscillatory rheology.

### Dynamic oscillatory rheology

We used a rotational rheometer DHR-3 (TA Instruments) to characterize the viscoelastic properties of resected human colonic mucus and HCDM. An amplitude sweep at a constant frequency of 1 rad/s was first performed using a 40 mm parallel plate geometry to determine the linear viscoelastic region (**Supplementary Methods SM1A**). Next, frequency sweeps (0.01 to 50–100 rad/s) were conducted at constant strain amplitude of 10% (0.5 μNm torque), with 5–10 measurement points per decade and a plate separation gap of 300 μm, to evaluate the loss modulus (G”) and storage modulus (G’). Mucus samples were characterized in DPBS containing calcium and magnesium (pH 7) at 25°C. No differences in the mechanical spectrum of mucus were observed between ambient (25°C) and body temperature (37°C) (**Supplementary Methods SM1B**).

### pH-dependent mechanical behavior of HCDM

Buffers at pH 2 and 9 were prepared by adjusting a phosphate buffer (pH 6) with sodium hydroxide and phosphoric acid, respectively. The pH was measured using a digital pH meter (VWR Symphony B10P), with the pH electrode calibrated before each use using standard calibration buffers. The HCDM samples were equilibrated in each buffer and then analyzed by dynamic oscillatory rheology as previously described.

### Proteolytic digestion

HCDM was submerged in a digestion solution at a 1:50 ratio of papain to glycoprotein in phosphate-buffered saline (PBS) supplemented with 2 mM L-cysteine to enhance papain’s enzymatic activity. The HCDM samples were then digested for 24 hours at 37°C under static conditions. Following treatment, the mechanical properties of both treated and untreated HCDM were determined by oscillatory rheology as described above. All experiments were performed in triplicate.

### Media preparation

Defined minimal media for cultivating *Bacteroides* species was prepared following the recipe by Varel and Bryant^90,91^, whereas chemically defined media for the cultivation of *Bifidobacteria* and *Enterococci* was prepared according to Socransky and coworkers^92^. Complex media used in this study included yeast casitone fatty acids (YCFA) with and without 0.5% glucose, peptone-yeast-glucose (PYG) broth, and supplemented brain heart infusion (BHIS). Agar plates were prepared by adding 1.5% agar to each broth. YCFA and PYG were prepared according to the DSMZ recipes, available on their website. BHIS were made using premixed formulations from Bacto™ Brain Heart Infusion (BD Diagnostics). The pH of all media formulations was adjusted to 6.8 - 7.0.

### Bacterial strains

The following bacterial strains were used: *Bacteroides fragilis* ATCC 25285 and *Bifidobacterium longum subsp. infantis* ATCC 15697 were obtained from the American Type Culture Collection (ATCC). *Bacteroides thetaiotaomicron* DSM 2079, *Bacteroides caccae* DSM 19024, *Faecalibacterium duncaniae* (previously named *Faecalibacterium prausnitzii*) DSM 17677, *Roseburia intestinalis* DSM 14610, *Ruminococcus gnavus* DSM 108212, and *Enterococcus faecalis* DSM 20478 (ATCC 19433) were purchased from the DSMZ-German Collection of Microorganisms and Cell Cultures. *B. fragilis, B. thetaiotaomicron, B. caccae, B. longum subsp. Infantis* and *E. facaelis* were routinely cultured anaerobically at 37°C in BHIS broth before being used in mucus utilization and degradation experiments. *F. duncaniae, R. intestinalis*, and *R. gnavus* were routinely cultured anaerobically at 37°C in YCFA. All bacteria were inoculated from frozen stocks stored at −80°C into 10 mL of their respective media overnight before being used with mucus utilization and degradation studies.

### Mucus utilization

We assessed HCDM utilization by measuring colony-forming units (CFUs) per mL using serial dilution and agar plating. *B. thetaiotaomicron*, *B. caccae* and *B. fragilis* were incubated for 24 h in minimally defined media, with and without ∼ 1 mg/mL HCDM, while *B. longum*, and *E. faecalis* were grown in chemically defined media under the same conditions. *F. duncaniae, R. intestinalis, and R. gnavus* were grown in glucose-free YCFA, with and without ∼ 1 mg/mL HCDM. The mucus concentration in the broth was in alignment with prior studies^46^. Agar plating was performed on BHIS agar for *B. fragilis*, *B. thetaiotaomicron*, *B. caccae*, *R. gnavus*, *B. longum, and E. faecalis,* and on YCFA agar for *F. duncaniae* and *R. intestinalis*.

### Mucus Degradation

Mucus degradation was evaluated using anaerobic media prepared two weeks in advance, ensuring minimal impact of cysteine on mucus structure (**Supplementary Methods SM2**). Cysteine, a reducing agent crucial for the growth of strict anaerobic bacteria, can reduce the disulfide bonds in mucus. After 36 hours of incubation—12 hours for mucus degassing and 24 hours for bacterial culture—the dynamic moduli of control mucus (without bacteria) decreased by approximately an order of magnitude but remained consistent across all media formulations used in this study. Including control mucus samples without bacteria in all experiments allowed us to effectively distinguish media-induced changes from bacterial degradation. Each experiment was conducted in 2 mL of broth using 5 mL culture tubes with closures (VWR 60818-565). For prolonged bacterial cultures lasting several days, 12-well plates with 0.4 µm Transwell inserts were used to allow daily media refreshment while maintaining a consistent broth volume (2 mL). Following incubation, mucus samples were collected and analyzed via oscillatory rheology.

### Enzymatic removal of mucin glycans

Mucin glycans were enzymatically removed using O-glycosidase, sialidase (α2-3,6,8,9 Neuraminidase A), and fucosidase (α1-3,4 Fucosidase), all purchased from New England Biolabs. O-glycosidase, which cleaves core 1 and core 3 O-linked disaccharides from glycoproteins, was used at a stock concentration of 40×10^6^ units/mL. Sialidase, with broad specificity for linear and branched non-reducing terminal sialic acid residues, had a stock concentration of 20×10^3^ units/mL, while fucosidase, which hydrolyzes terminal α1-3 and α1-4 linked fucose residues, was obtained at a stock concentration of 4×10^3^ units/mL. For glycan removal, 100 µL mucus were incubated with either 10 or 100 µL/mL of each enzyme at 37°C for 1 hour. All reactions were performed using the manufacturer’s recommended buffers and protocols available on their website.

### Protease inhibitor assay

To evaluate the role of proteases, *B. fragilis* was grown for 24 hours in 10 mL of defined minimal media containing 0.5% glucose, while *B. longum* and *E. faecalis* were cultured in glucose-free YCFA under the same conditions. Following incubation, culture supernatants were collected and filter-sterilized using a 0.2 μm filter. A protease inhibitor cocktail (Sigma-Aldrich, catalog #SIAL-P8465-5ML), effective against serine, cysteine, aspartic, and metalloproteases as well as aminopeptidases, was prepared by reconstituting the lyophilized powder with 1 mL of DMSO and 4 mL of deionized water, according to the manufacturer’s instructions. Sterile mucus (∼1 mg/mL) was incubated with filter-sterilized culture supernatants, either with or without the protease inhibitor cocktail, at 37°C for 12–24 hours. Mucus viscoelasticity was subsequently assessed using oscillatory rheology. Control samples consisted of sterile mucus incubated in culture media under identical conditions.

### Crystal violet staining

*E. faecalis* was grown overnight anaerobically at 37°C. Cultures were diluted to an OD600 of 0.05 in 2 mL of YCFA media, and mucus was deposited in Transwell inserts (12-well plates, 2 mL volume) and degassed overnight. Three uninfected blanks and three infected wells were prepared. After infection, plates were incubated anaerobically at 37°C for 48 hours. Following incubation, Transwell inserts were washed in sterile PBS to minimize biofilm disturbance, dried at room temperature for 10–15 minutes, and stained with 0.1% (w/v) crystal violet for 10 minutes. Excess stain was removed, and wells were washed with ultrapure water until clear. Membranes were excised from the Transwell inserts and transferred to microcentrifuge tubes. Crystal violet was solubilized with 500 µL of 30% acetic acid per well for 10–15 minutes at room temperature. The solution was mixed, and 300 µL was transferred to a 96-well plate. Serial 1:2 dilutions were prepared with 30% acetic acid, which also served as the blank. Optical density was measured at 570 nm to quantify biofilm-associated crystal violet.

### Lectin staining of exopolysaccharides

Wheat germ agglutinin (WGA) conjugated to Alexa Fluor 488 was used to visualize mucus fibers, and concanavalin A (ConA) labeled with Texas Red was used for exopolysaccharide staining. Mucus samples infected with *E. faecalis* for 48 hours, were placed on poly-L-lysine-treated glass slides and air-dried for 30 minutes at room temperature. Samples were fixed with 4% formaldehyde at 37°C for 15 minutes, followed by three 10-minute washes with PBS at room temperature. Slides were incubated in the dark with 50 µL of 5 µg/mL WGA in PBS for 10 minutes, then washed twice with PBS. They were subsequently stained with 50 µg/mL ConA for 30 minutes at room temperature, followed by three 5-minute PBS washes. Samples were counterstained with ProLong Gold Antifade with DAPI to visualize nuclei and stored at −20°C until imaging.

### RNA-Sequencing and differential gene expression analysis

RNA was extracted from *E. faecalis* using the RNeasy PowerMicrobiome Kit (Qiagen) according to the manufacturer’s instructions. Bacterial cultures were incubated for 12 h in YCFA basal broth (pH 6.8-7) under four conditions, each in triplicate: no sugars, mucus (approx. 1 mg/mL), mucus + 0.5% glucose, and 0.5% glucose. These conditions were tested under both anaerobic and aerobic environments. Following RNA extraction, fragment analysis was performed to assess RNA integrity. Novogene Corporation carried out sample quality control, library preparation, and sequencing. The quality of sequencing data was evaluated using FastQC. Sequencing reads were aligned to the *E. faecalis* ATCC 19433 reference genome using Bowtie2. Post-alignment processing, including conversion, sorting, and indexing of alignment files, was done with SAMtools. Gene expression was quantified using featureCounts, and the resulting count files were imported into R for further analysis. Differential gene expression analysis was conducted using the DESeq2 package. Differential expression was determined using a significance threshold of adjusted *p-value* (padj) < 0.05 and an absolute log2 fold change (|log2FC|) > 1. Sequences of differentially expressed genes were used to identify homologous sequences in other organisms, obtaining UniRef90 identifiers. These UniRef90 identifiers were sorted and filtered based on the best sequence matches. Functional annotations for these identifiers, including protein functions, pathways, and gene ontology (GO) terms, were retrieved from the UniProt database. Enrichment analysis of GO terms was performed using Fisher’s Exact Test, with *p-*values calculated relative to the *E. faecalis* background genome. GO terms linked to transcription, translation, DNA replication, cell cycle regulation, ribosomal processes, and nucleotide metabolism were excluded. Multiple testing corrections were applied using the Benjamini-Hochberg method.

### Statistical analysis

Data are presented as mean ± standard deviation. Pairwise comparisons were performed using the t-test with a two-tailed distribution, with statistical significance defined as *p* < 0.05 (*) or *p* < 0.01 (**). For comparisons across multiple conditions, one-way ANOVA was employed. Significant differences were further analyzed using Tukey’s post-hoc test at a significance level of 0.05.

## Acknowledgments

The authors acknowledge the use of facilities and instrumentation supported by the NSF through the Cornell University Materials Research Science and Engineering Center (DMR-1719875). The authors also thank Héctor Loyola Irizarry, Nandika Nair, and Carl Soderstrom for their technical support throughout this project. This work was funded in part by a grant from the Herbert W. Hoover Foundation. E.V.W received funding from the Natural Sciences and Engineering Research Council of Canada (NSERC) through the Postgraduate Scholarships – Doctoral (PGSD) program.

