## Supplementary Information for "Environmental factors drive bacterial degradation of gastrointestinal mucus"

**Contents:**

**Supplementary Figures** S1 to S9

**Supplementary Methods** SM1 to SM2

### Supplementary Figures

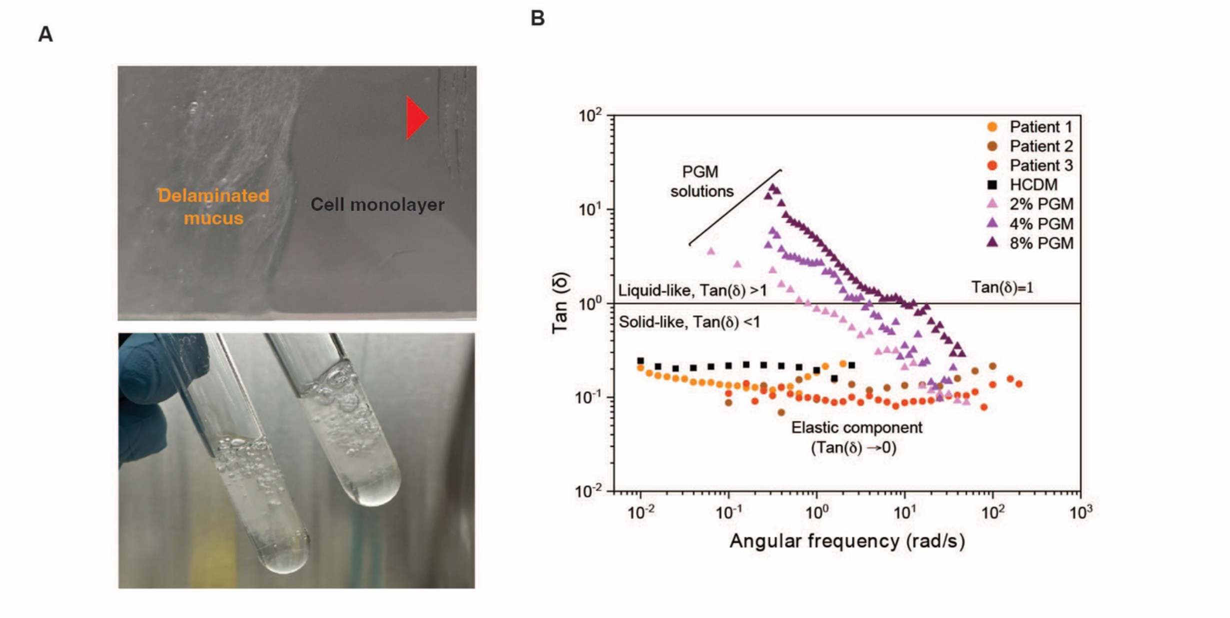


**Supplementary Figure S1. Evidence of mucus delamination and similarity to mucus derived from human biopsies. (A)** Delaminated mucus after treatment with 1 mM forskolin. The red arrow indicates a scratch made on the flask to demonstrate that the cell monolayer is left intact. About 1 mL of hydrated mucus is collected per T-25 flask. **(B)** Plot of the phase angle for human colonic biopsies, HCDM, and porcine gastric mucin (PGM). The phase angle describes the ratio of viscous to elastic effects and is a sensitive indicator of cross-linking. Values close to zero across a wide frequency range demonstrate the network structure in HCDM and human mucus, responsible for their viscoelastic behavior. In contrast, uncross-linked polymers like PGM exhibit a phase angle inversely proportional to the frequency.

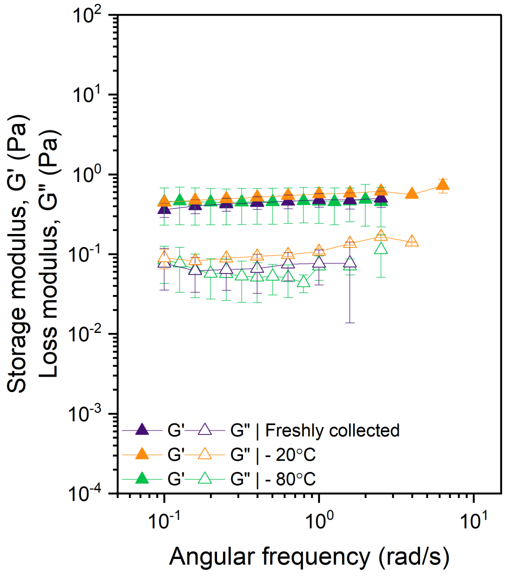


**Figure S2. Impact of freezing on HCDM mechanical properties.** The mechanical properties of HCDM were assessed after freezing at -20°C or -80°C for 1 week and subsequently thawing to room temperature.

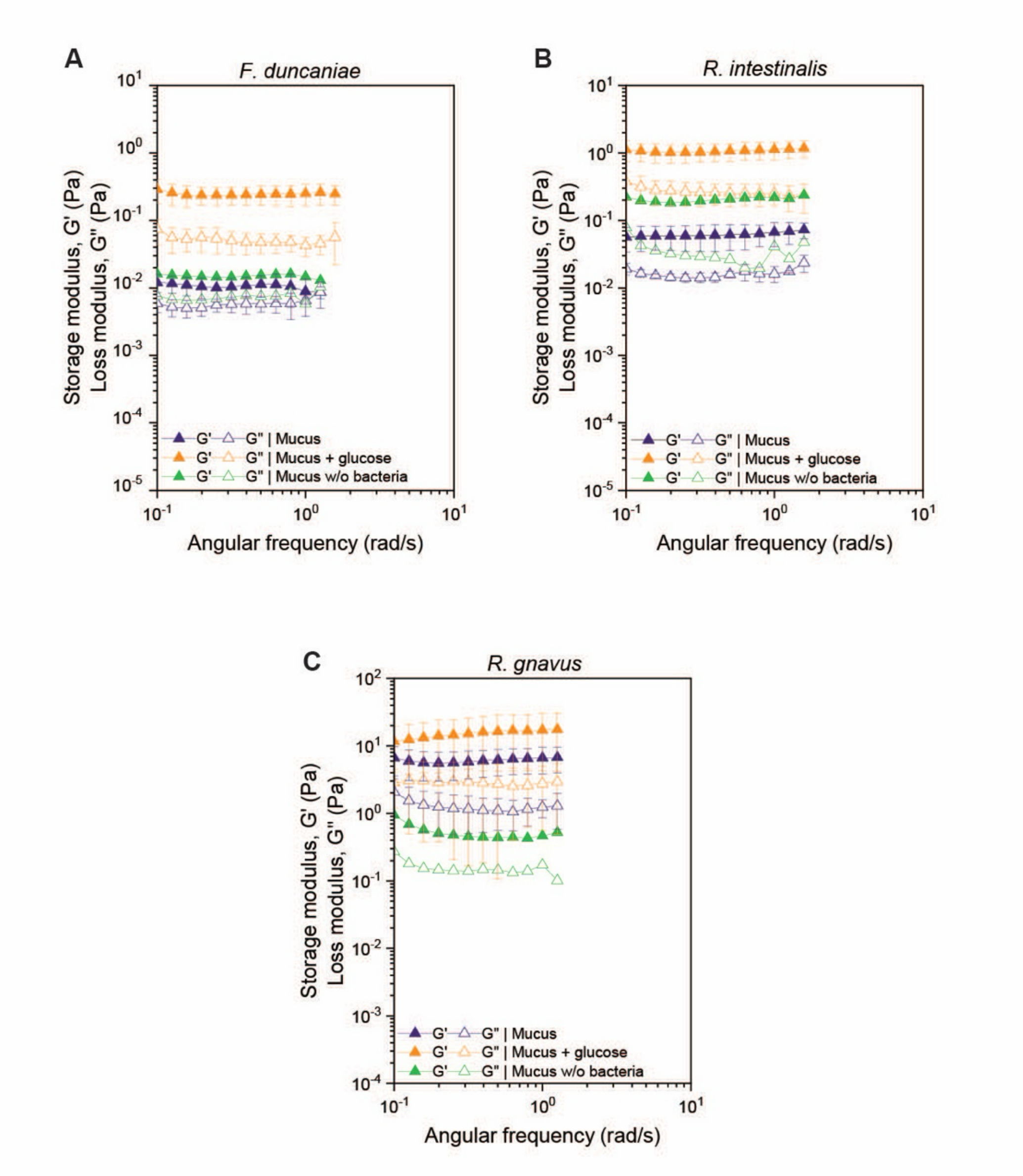


**Supplementary Figure S3. Mucus bulk rheology remained unchanged after co-culture with select gut microbiota.** Oscillatory rheology was performed on mucus cultured with **(A)** *F. duncaniae,* **(B)** *R. intestinalis*, and **(C)** *R. gnavus* with or without glucose. Mucus incubated without bacteria under similar conditions served as the control. Curves represent the average of three biological replicates.

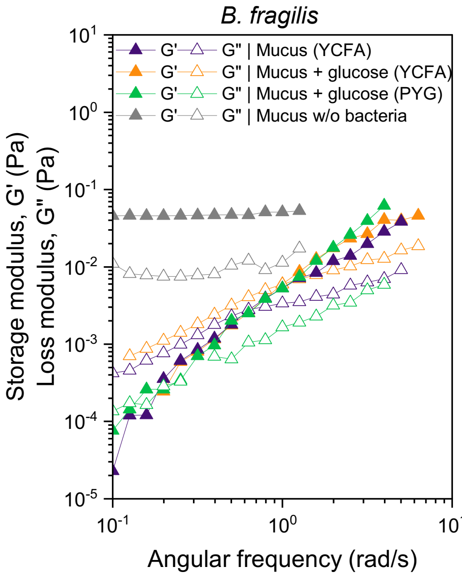


**Supplementary Figure S4. *B. fragilis* degrades intestinal mucus under nutrient-rich conditions.** Oscilatory rheology performed on mucus co-incubated with *B. fragilis* in glucose-free YCFA media, YCFA media supplemented with glucose, or PYG media with glucose.

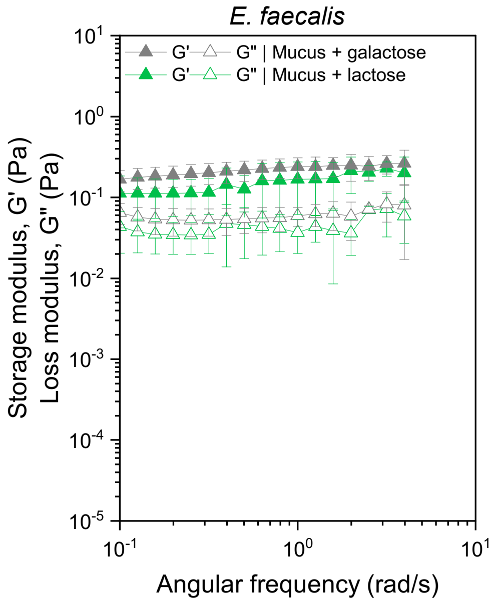


**Supplementary Figure S5. Mucus degradation by *E. faecalis* is suppressed by simple sugars.** Oscillatory rheology performed on mucus incubated with *E. faecalis* in the presence of galactose or lactose. Data corresponds to the average of 3 different measurements, collected at 24 hours incubation period in anaerobic conditions

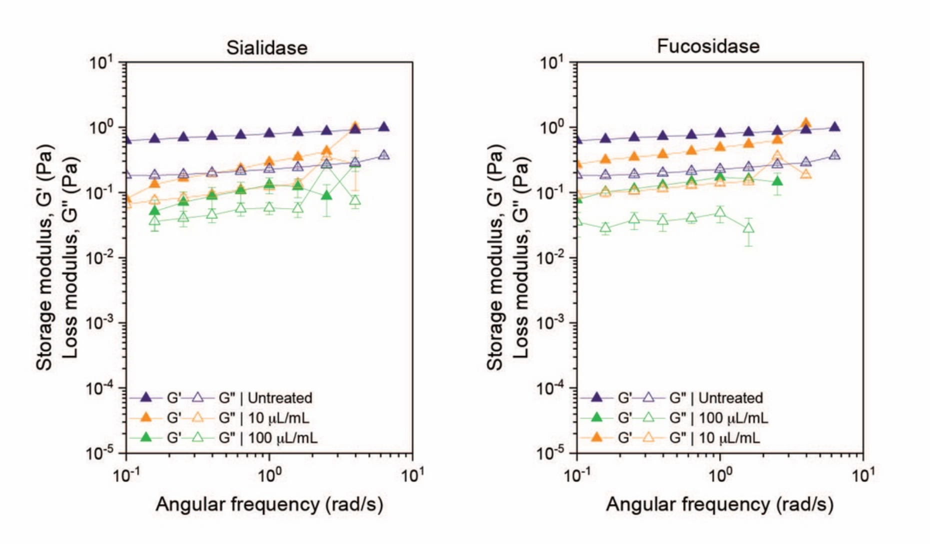


**Supplementary Figure S6. Impact of Enzymatic Glycan Removal on HCDM Mechanical Properties.** Oscillatory rheology was performed on mucus after incubation with sialidase (left) and fucosidase (right) or without any addition of enzymes (untreated).

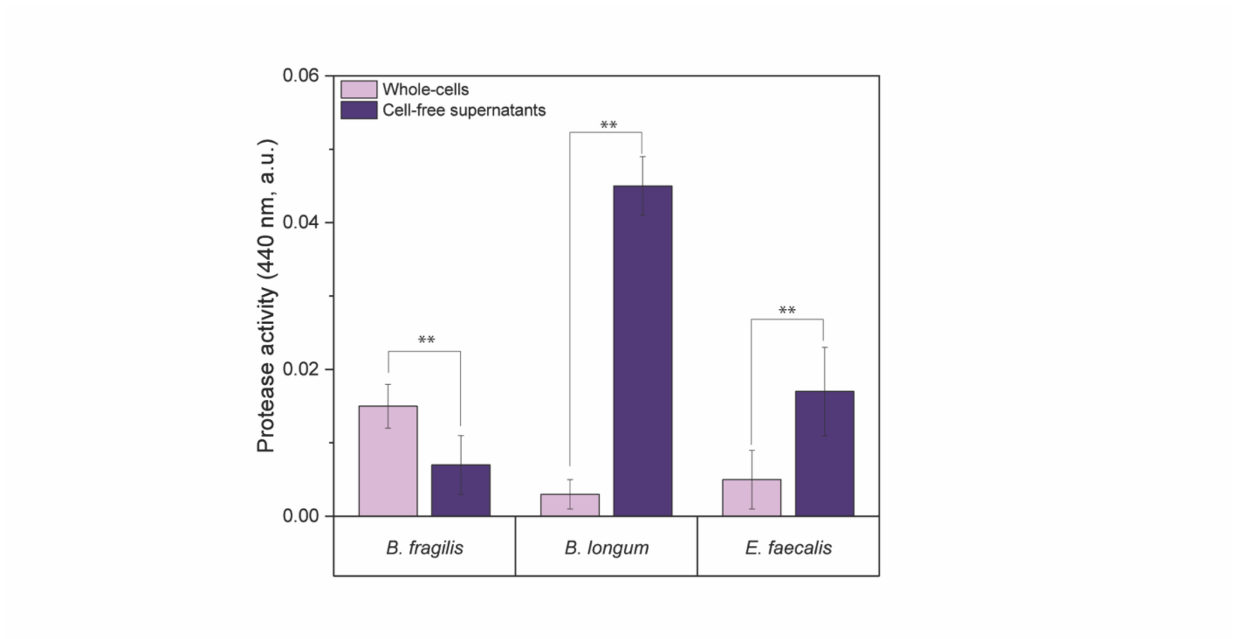


**Supplementary Figure S7. Protease activity of *B. fragilis*, *B. longum*, and *E. faecalis* measured using azocasein as a substrate.** Protease activity was measured in both whole cells and cell-free supernatants. *B. fragilis* was cultured in defined minimal media with 0.5% glucose, whereas *B. longum* and *E. faecalis* were cultured in glucose-free YCFA. Data was normalized to the blank (azocasein without proteases). Bars represent mean ± SD. Asterisks denote statistical significance: $p < 0.01$ (**).

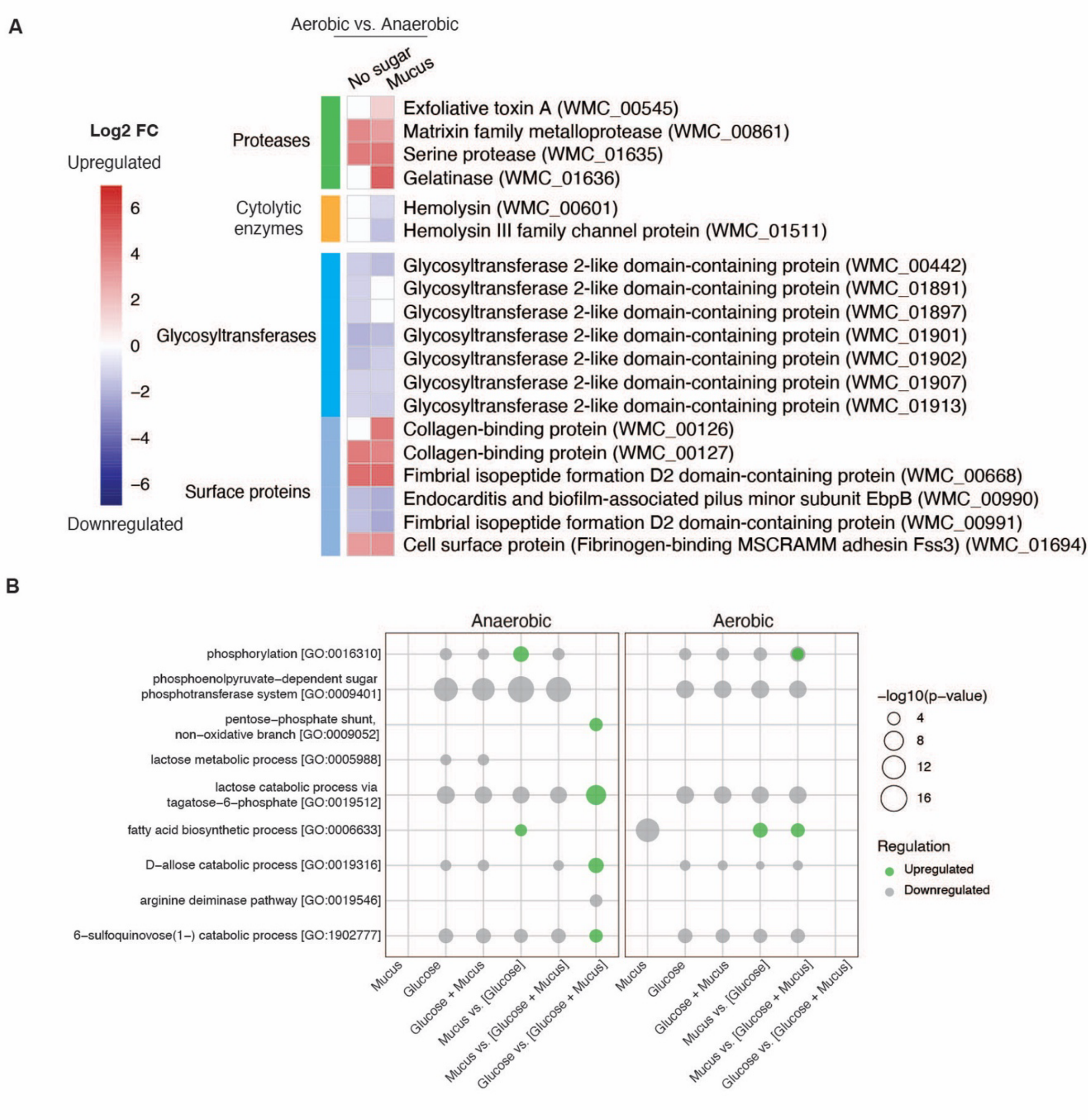


**Supplementary Figure S8. Metabolic Pathway Enrichment and Virulence Gene Expression in *E. faecalis*. (A)** Significantly enriched metabolic pathways in *E. faecalis* under anerobic and aerobic conditions. **(B)** Genes involved in *E. faecalis* virulence were selected for heatmap. Differential gene expression analysis was conducted using DESeq2 with an adjusted *p-value* threshold of <0.05 and a $|\log_2 \text{FoldChange}| > 1$.

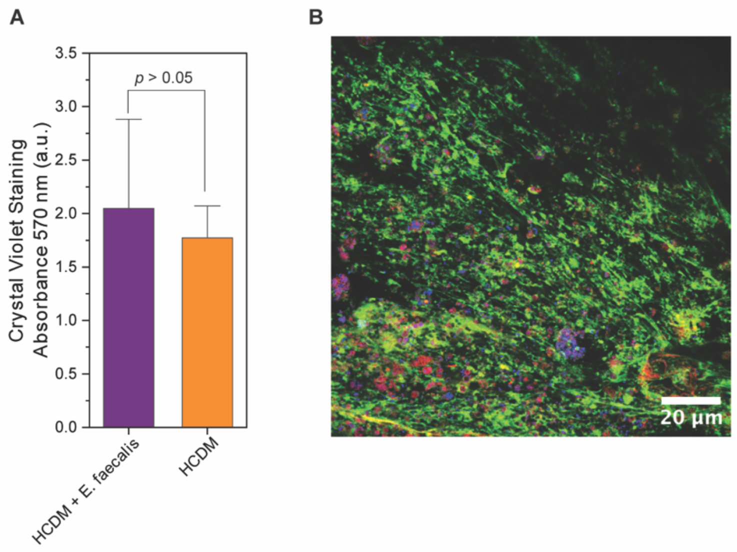


**Supplementary Figure S9. Assays to assess biofilm formation in *E. faecalis.* (A)** Crystal violet staining of *E. faecalis* after 48 hours of anaerobic culture on mucus with glucose. **(B)** Visualization of the mucus and extracellular polymeric substances using wheat germ agglutinin (WGA) to stain mucus (green), concanavalin A (ConA) to label bacterial exopolysaccharides (red), and DAPI to stain bacterial cells (blue). Bars represent mean ± SD. Statistical significance: $p < 0.05$

### Supplementary Methods

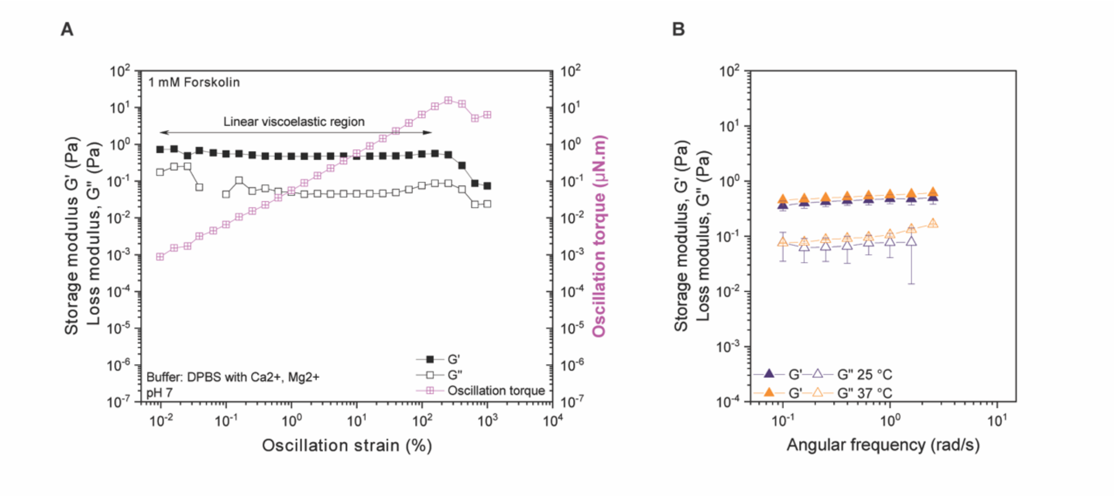


**Supplementary Methods SM1. Mechanical and Rheological Characterization of Human Cell-Derived Mucus (HCDM). (A)** Linear viscoelastic region (LVR) for human cell-derived mucus (HCDM) determined using an amplitude sweep at a constant frequency of 1 rad/s. **(B)** The mechanical spectrum of HCDM was analyzed at both ambient temperature (25°C) and body temperature (37°C).

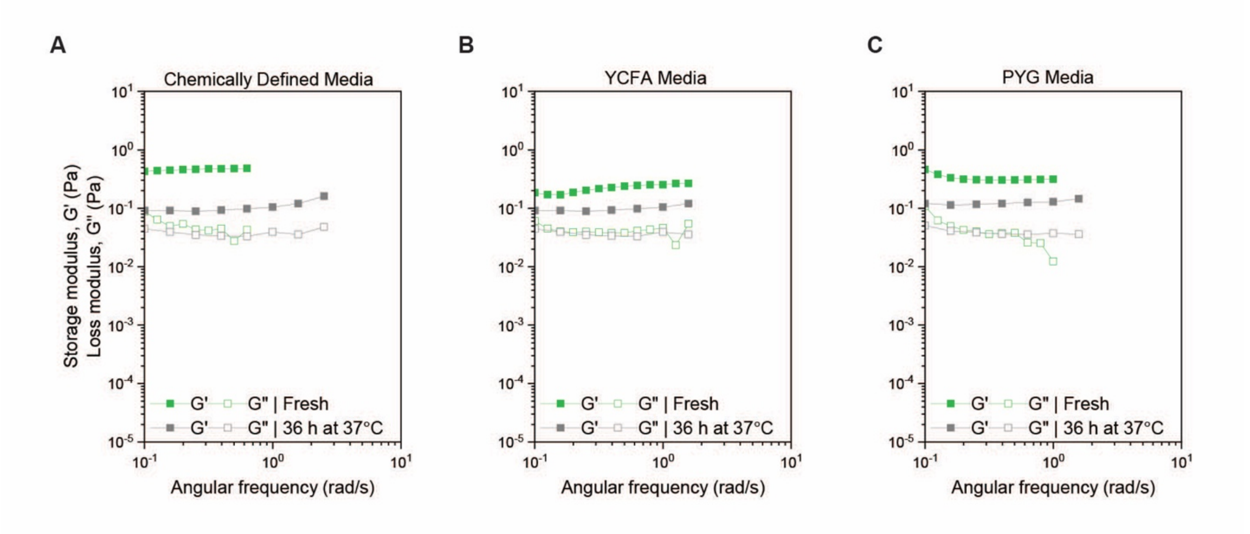


**Supplementary Methods SM2. Effect of cysteine-containing anaerobic broths on HCDM rheological properties before and after incubation for 36 hours at 37°C.** Results are shown for (A) chemically defined media, (B) yeast-casitone-fatty acids (YCFA) media, and (C) peptone-yeast-glucose (PYG) media. Cysteine is a reducing agent essential for the cultivation of strict anaerobes that can affect the rheological properties of intestinal mucus.
